# Intrahippocampal delivery of hyperphosphorylated human tau oligomers induces neurodegeneration in non-transgenic wildtype mice

**DOI:** 10.1101/2025.03.10.642468

**Authors:** Hong-Ru Chen, Hsiao-Tien Hagar, Kuan-Wei Wang, Stacy Hovde, Hsiang-Ying Lin, Min-Hsuan Lin, Cheng-Chun Chang, Ching-Yung Wang, Chia-Yen Chen, Emily J Bian, Melissa M. Kinkaid, Min-Hao Kuo, Chia-Yi Kuan

**Affiliations:** Department of Neuroscience, University of Virginia School of Medicine, Charlottesville, VA, USA; Department of Biochemistry and Molecular Biology, Michigan State University, East Lansing, MI, USA; Department of Life Sciences and Institute of Genome Sciences, National Yang Ming Chiao Tung University, Taipei, Taiwan; Brain Research Center, National Yang Ming Chiao Tung University, Taipei, Taiwan

**Author notes:** Correspondence: Chia-Yi Kuan Department of Neuroscience, University of Virginia School of Medicine, 409 Lane Road, MR-4, 4046, Charlottesville, VA 22908, USA Min-Hao Kuo Department of Biochemistry and Molecular Biology, Michigan State University, 603 Wilson Road, Room 401, East Lansing, MI 48824, USA.

## Abstract

Hyperphosphorylated tau (p-tau) forms neurofibrillary tangles, a key biomarker for Alzheimer’s disease and additional neurodegenerative tauopathies. However, neurofibrillary tangles are not sufficient to cause neuronal dysfunction or death. Intrahippocampal injection of tau isolated from AD patients has limited effects on the cognitive functions of non-transgenic mice, despite the recapitulation of pathological tau deposits in the mouse brain. It therefore remains uncertain as to whether all hyperphosphorylated tau is directly responsible for AD neurodegeneration. We examined this issue by injecting recombinant p-tau oligomers to the hippocampus of non-transgenic, wildtype mice and found progressive cognitive deficits that correlate with neuron death spreading from the ipsilateral hippocampus to the cortex. Apomorphine, which retards p-tau aggregation and cytotoxicity *in vitro*, antagonized p-tau-induced cognitive deficits and neuron death. These results suggest the pathogenic role of p-tau oligomers and a novel AD model facilitating drug development.

## INTRODUCTION

Hyperphosphorylated tau (p-tau) is the major component of neurofibrillary tangles (NFTs) in the brain of Alzheimer’s disease (AD) patients (*1, 2*). Several lines of evidence suggest that p-tau oligomers may be the critical driver of neurotoxicity and disease progression. Firstly, the spatiotemporal distribution of p-tau and NFTs, but not the amyloid plaques, correlates with the progression of cognitive decline in AD patients, as shown in Braak-neuropathological staging (*3*). Cell culture and transgenic mouse models demonstrated that p-tau is required for the deleterious effects of amyloid-β peptides (*4, 5*), consistent with numerous reports that neurodegenerative tauopathies, except AD, are associated with p-tau deposits without significant amyloid plaques (*6, 7*). Furthermore, several anti-amyloid treatments are potent in reducing the amyloid burden, but only show minimal benefits in preserving cognitive functions (*8–10*).

Many current transgenic tauopathy mouse models express mutant tau alleles, but the majority of late-onset AD involves hyperphosphorylated p-tau oligomers and tangles without specific tau mutations (*11*). Intrahippocampal injection of tau extracted from the AD and tauopathy patient brains to mice recapitulated tau histopathology, but few produced behavioral or cognitive deficits if wildtype mice were injected (*12–15*). Similar results were reported with the use of humanized mice in which both copies of the rodent tau gene, MAPT, were replaced with the human orthologue (*16*). This disparity between cognitive integrity and conspicuous neurofibrillary tangles suggests that the common protocols of tau isolation from AD patient brains may contain an inadequate amount of the pathogenic tau species to induce neurodegeneration. Thus, introducing a tau species with the pathogenic activity may improve animal modeling in the revelation of AD pathogenesis and in drug development.

To this end, we use hyperphosphorylated tau produced by the PIMAX system. In PIMAX (Protein Interaction Module-Assisted function X), tau and one of its main kinases, GSK-3β, are each expressed in *E. coli* as a chimeric protein fused to the leucine zipper domain of Jun and Fos, respectively (*17*). Fos-Jun heterodimerization brings tau and GSK-3β to close proximity, resulting in highly efficient phosphorylation of tau (*18*). P-tau obtained from PIMAX possesses molecular traits highly relevant to the disease, including disease-related hyperphosphorylation marks, inducer-free aggregation, and cytotoxicity to the SH-SY5Y and HEK293T cells (*18–20*). In addition to impairing the viability of these cells, p-tau is free of lipopolysaccharides and activates microglial cells induced from human pluripotent cells (*25*), a feature that is consistent with the increasing recognition of the involvement of neuroinflammation and microglia in neurodegeneration.

Besides mechanistic studies, p-tau from the PIMAX expression was also applied to AD drug discovery endeavors. Apomorphine was one of the top hits emerging from a pilot drug screen for compounds with significant inhibitory activities against the formation of cytotoxic p-tau aggregates in vitro (*19, 20*). Best known as a dopamine receptor agonist being used to treat the off episodes of Parkinson’s disease, apomorphine enhanced short-term memory functions in 3xTg-AD mice, although the mechanism was attributed to promoting amyloid-β degradation (*21*). In vitro, apomorphine exhibited highly effective antagonizing functions against p-tau induced ER stress and Unfolded Protein Response, accumulation of reactive oxygen species and cytoplasmic calcium, apoptosis, and cell death (*20*). Intrigued by the functions of apomorphine in suppressing p-tau toxicity and preserving memory in 3xTg-AD mice, we posited that neurological deficits caused by p-tau could be ameliorated by apomorphine treatment. If proven, p-tau and apomorphine might be used to establish a p-tau-induced preclinical neurodegenerative model as an alternative to transgenic mutant tau mice for drug discovery, considering that tau mutations are rare in Alzheimer’s disease patients.

## RESULTS

### Hyperphosphorylated human tau oligomers induce mitochondrial superoxide accumulation and neurite fragmentation

To evaluate the pathogenic roles of p-tau, we first examined how p-tau affected the integrity of primary neurons in vitro. The p-tau used in this study (1N4R) existed primarily as SDS-sensitive oligomers between 66 and 440 kDa in size, as shown by size exclusion chromatography (Fig. 1A). Mouse cortical neurons were isolated from E16 embryos, incubated for 6 days *in vitro* (DIV), and then treated with 6 µM of recombinant non-phosphorylated or p-tau prepared in parallel. We first focused our attention on the mitochondria of these primary neurons, since mitochondrial dysfunction is well-recognized as an important driver of neuronal loss and neurodegeneration. MitoSOX and MitoTracker dyes were used to examine the mitochondrial superoxide levels and morphology two days after the addition of tau or p-tau, when there were no discernible changes in the number or the morphology of cells. Strong mitochondrial superoxide signals (MitoSOX) were seen in neurons treated with p-tau but not with the unmodified tau (Fig. 1B), indicating significant accumulation of reactive oxygen species. The levels of such radicals appeared to be positively correlated with the dose of p-tau, consistent with our previous findings with SH-SY5Y and HEK293T cells (*18–20*). To further explore the consequences of p-tau treatment, MitoTracker was used to visualize the shape and distribution of mitochondria. Figure 1C shows condensation of mitochondria in p-tau-, but not tau- or PBS mock-treated sample. Besides increasing the abundance of mitochondria in the soma, p-tau also changed the mitochondrial morphology from long and tubular to a round shape. These mitochondria were also stained strongly with MitoSOX (Fig. 1C; arrow head), suggesting a dysregulated redox status following p-tau treatment. Tau is a microtubule-binding protein; hyperphosphorylation squelches the affinity of tau for microtubules. One possible mode of action for cytotoxic p-tau is to recruit and interfere with the function of the endogenous tau, leading to cytoskeleton disintegration in cells (*20*). We were curious as to whether p-tau treatment resulted in alterations in neurites of primary neurons. To this end, we probed neurites with anti-MAP2 antibodies, and found that four days after the addition of the recombinant protein, p-tau dose-dependent massive fragmentation of neurites was observed (Fig. 1B). The unmodified tau did not have this effect. The detrimental effects of p-tau on neurites can be explained by the reduction of relevant proteins. For example, p-tau caused significant reduction of MAT2, and α- and βIII-tubulins (Sup. Fig. 1A-C). In contrast, a moderate increase was seen in the activated Caspase 3 (Sup. Fig 1B, C). Unsurprisingly, the prominent disorganization of neurites was linked to neuronal viability loss, as evidenced by cellular viability tests of propidium iodide (PI) uptake and tetrazolium colorimetry assays (Sup. Fig. 1D). These toxic responses were either absent or were much less severe when tau was used, in accordance with our previous reports of p-tau-specific cytotoxicity to SH-SY5Y and HEK293T cells (*19, 20*). Lastly, the neuronal import of the recombinant tau and p-tau from the medium was confirmed by the use of FLAG-tagged p-tau and tau for treatment. Immunoblots revealed the presence these two FLAG-tagged proteins in cell lysates (Fig. 1D).

**Fig. 1.**
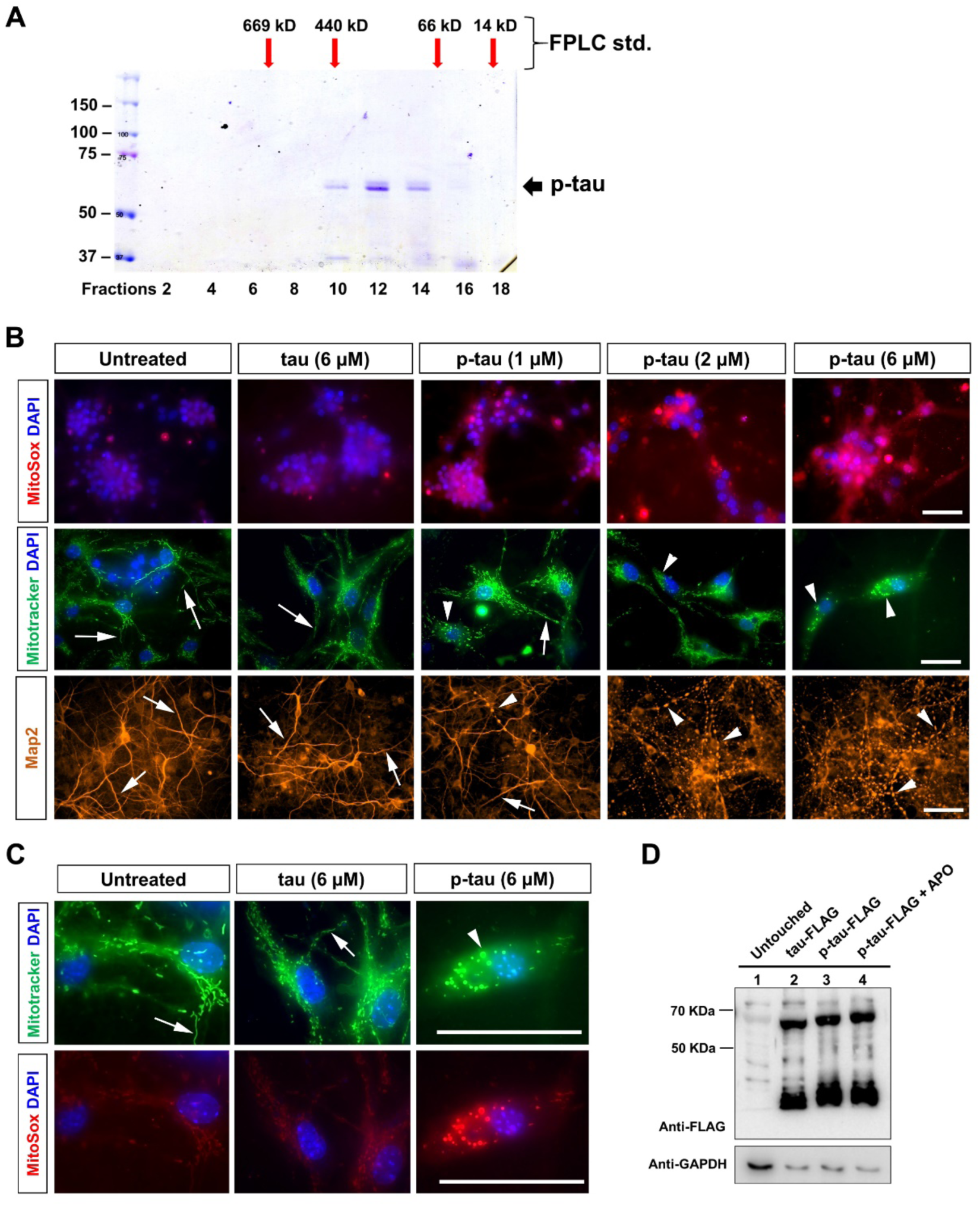
Hyperphosphorylated human tau oligomers induces mitochondrial superoxide production and neurite fragmentation while non-phosphorylated tau exhibits no such effects. **(A)** Gel filtration analysis revealed that hyperphosphorylated tau (p-tau) produced by the PIMAX system predominantly exists as ∼250 kD oligomers. The elution fractions of molecular weight standards (std) are shown above the gel image. **(B-D)** Cortical neurons isolated from embryonic day 16.5 (E16.5) C57BL/6 mice were cultured in vitro for 6 days, followed by a 2- or 4-day treatment with varying concentrations of non-phosphorylated tau or p-tau oligomers. **(B)** Mouse cortical neurons were treated with unphosphorylated tau or p-tau at the indicated concentrations for 2 days, followed by MitoSox and Mitotracker staining (top and middle panels). The arrows indicate normal, long tubular mitochondrial morphology, while the arrowheads point to short, fragmented mitochondria accumulated in the soma (middle panels). Neurites of cortical neurons showed p-tau dose-dependent fragmentation (lower panel). Neurites were visualized by immunofluorescence staining with anti-Map2 antibodies on neurons treated with p-tau or tau for 4 days. Clear fragmentation is seen in p-tau, but not in tau-treated neurons. The arrows indicate normal neurites, while the arrowheads point to fragmented neurites. Representative results from six independent cultures are shown. Scale bars, 50 µm. **(C)** High-magnification view of the specified group from Panel (B). Scale bars, 50 µm. **(D)** Uptake of tau and p-tau by primary neurons. FLAG-tagged tau and p-tau were used to treat cortical neurons for 4 days before lysate preparation and immunodetection with anti-FLAG antibodies. (n = 3).

### Spreading of tauopathy occurs in the WT mouse brain following hippocampal injection of hyperphosphorylated tau

After confirming the toxicity of p-tau to neurons, we asked whether this molecular outcome of p-tau treatment would translate into neurodegeneration if p-tau was delivered to the hippocampus stereotactically. We used the stereotaxic coordinates for the left dorsal hippocampus (*13, 22*) of 2- or 3-month-old wildtype, non-transgenic mice to introduce 2 µg of p-tau, unmodified tau, or PBS vehicle control (all 2 µl in volume) (Fig. 2A). A batch of mice were euthanized 2 days after the procedure to verify the injection location. P-tau produced by PIMAX was shown previously to display both AT8 and MC1 epitopes that represent hyperphosphorylation and the paired helical filament (PHF) conformation, respectively (*17, 18, 23, 24*). We conducted immunohistochemical staining of mouse brain sections with AT8 and MC1 antibodies and observed ipsilateral side-specific signals of AT8 and MC1 in the dentate gyrus (DG) and the hilus region of p-tau-injected mice (Fig. 2B). In contrast, the contralateral hippocampus lacked these immunosignals (Fig. 2B). We also stained these sections for Iba1, a marker for activated macrophage/microglia, as a test for the defensive immune response against foreign molecules. At the ipsilateral side of the p-tau group, Iba1 signals were easily noticed (panel h, Fig. 2C). Frequent overlaps between AT8 and Iba1 signals were found (arrows, panels h and i, Figure 2C), suggesting engulfment of the p-tau molecules by these immune cells. The unphosphorylated tau was probed with the phosphorylation-independent anti-tau DA9 antibodies, which also showed positive detection at the ipsilateral hippocampus (panels k and l, Fig. 2C). Some of the DA9 and Iba1 signals overlapped (arrows). These results suggested that the incoming recombinant proteins induced immune responses within two days after the injection. Compared with the p-tau contralateral side, or the tau ipsilateral DG regions, p-tau attracted significantly more Iba1 marked cells (compare panel h to k, Figure 2C), an observation reminiscent of the report that endotoxin-free p-tau activated microglial cells (*25*). One month after the injection, the AT8 epitope of tau hyperphosphorylation spread from the initial DG region to the CA1 (cornu ammonis) subfield of hippocampus, and to lesser extents to CA2 and CA3 (Fig. 3A-C; Sup. Fig. 2 for additional micrographs). Of note is the presence of AT8 signals at the anterior cingulate (AC) cortex. Importantly, AT8 signals also extended to the contralateral side, both AC cortex and the hippocampus fimbria region (Sup. Fig. 2A). Figure 3C illustrates the distribution of AT8 signals one month after p-tau injection. We further examined the pathological tau PHF conformation with the MC1 antibodies and found a similar expansion of this tau pathology (Figure 3D-F). However, the MC1 signals at the contralateral AC cortex and hippocampus appeared to be weaker than those for AT8, suggesting that tau hyperphosphorylation spread faster than did the PHF conformation. These results demonstrated the spread of abnormal phosphorylation and conformation of tau from the injection site to the distal, anatomically connected regions (*12–15*), in agreement with the prion-like p-tau transmission after injection of patient tau (*12, 13, 15*).

**Fig. 2.**
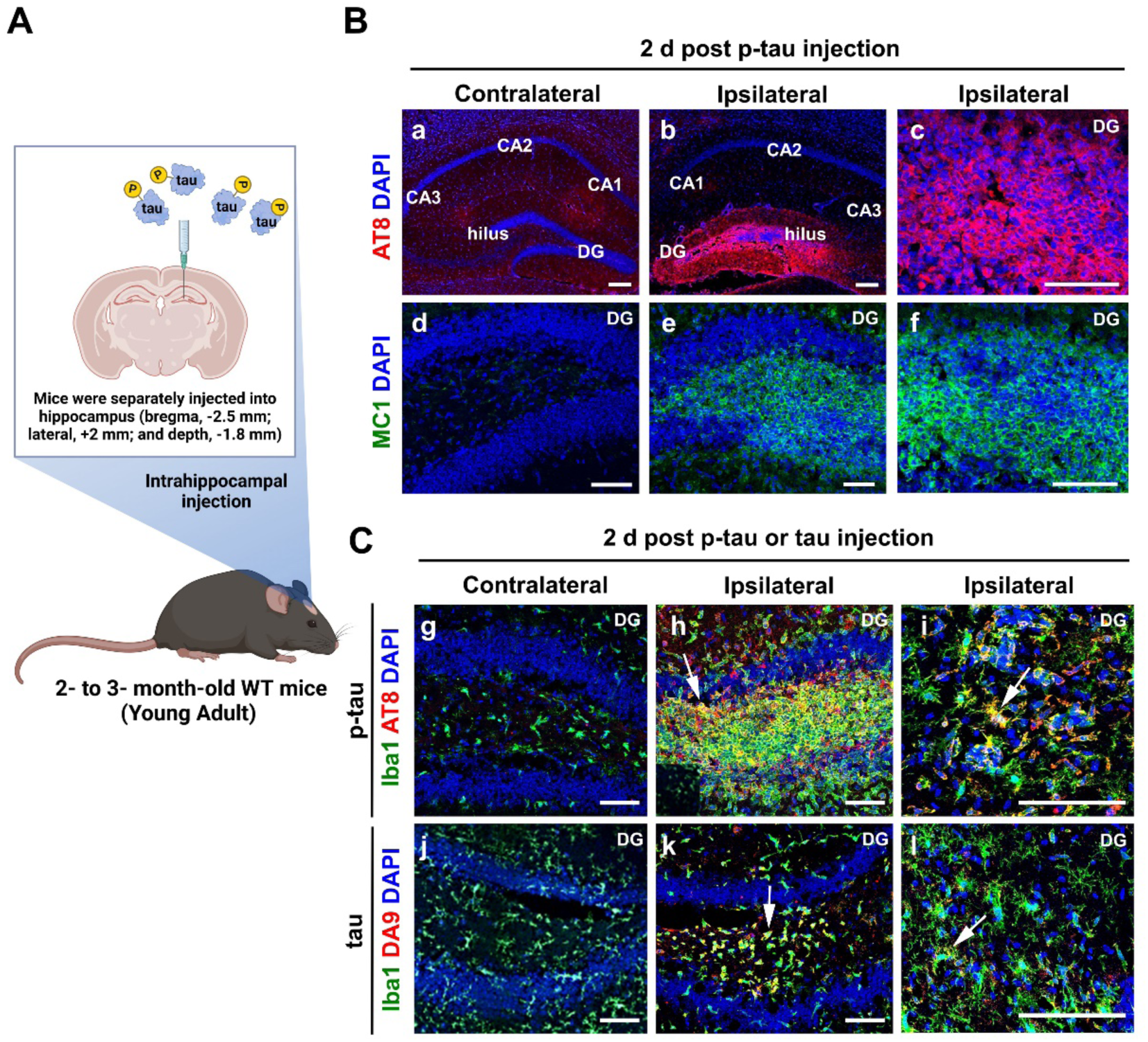
Delivery of hyperphosphorylated human tau oligomers and non-phosphorylated tau into the hippocampus of WT mouse brains following injection. **(A)** Stereotaxic coordinates for intrahippocampal injection of p-tau in this study. **(B)** Immunostaining conducted on 2-day post-injection mice brain sections suggested the successful delivery of p-tau to the ipsilateral hippocampus. AT8 (a-c) and MC1 (d-f) antibodies detected the presence of these epitopes at the ipsilateral dente gyrus and hilus region. DG, dentate gyrus. Scale bars, 50 µm. **(C)** Anti-Iba1 antibodies detected the presence of activated microglia/macrophages at or near the site of p-tau injection (h, i). Overlapping between Iba1 and AT8 fluorescence was frequently observed (arrows). DA9^+^ (g-i) pan-tau staining at 2 d post non-phosphorylated tau injection. Almost all recombinant tau was engulfed by Iba1^+^ microglia 2 days post-injection (arrows). Scale bars, 50 µm. (n = 3-5 for each condition).

**Fig. 3.**
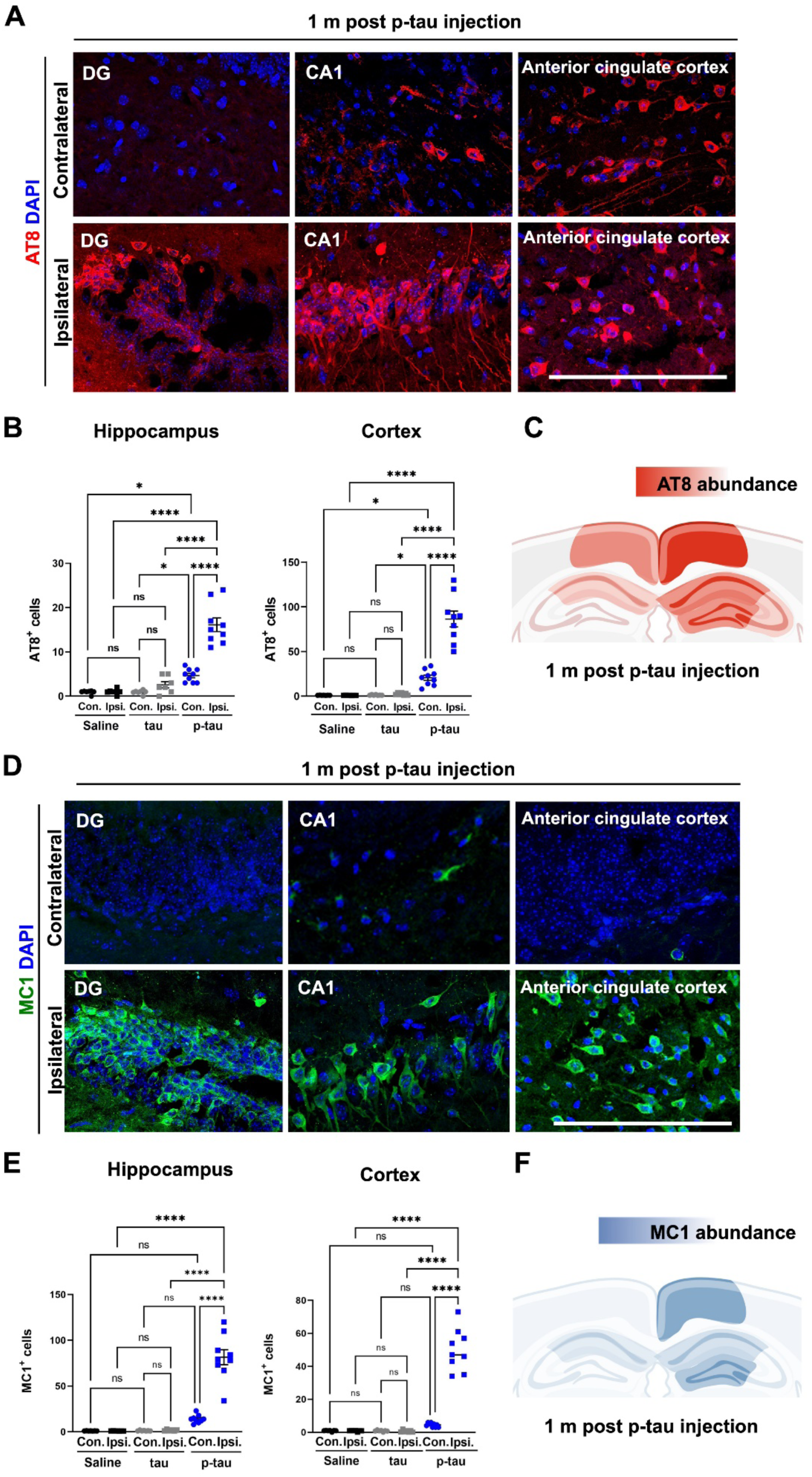
Transmission and spreading of tauopathy in the WT mouse brain following hippocampal injection of hyperphosphorylated tau. **(A)** One month after the p-tau injection, AT8 signals had spread to both the ipsilateral and contralateral CA1 sectors, as well as the anterior cingulate cortex. Scale bars, 50 µm. **(B)** Quantification of AT8^+^ p-tau protein in different areas of brains from saline, non-phosphorylated tau, and p-tau injection groups. Shown are means ± SEM. P-vale was determined by one-way ANOVA with Tukey’s multiple comparison post-hoc test. (*P<0.05, ***p=0.0001, ****p<0.0001). **(C)** Schematic depiction of AT8 signal seeding and spreading in the hippocampus and cortex. **(D)** Tau fibrils, as indicated by MC1 antibody reactivity, were observed in the contralateral and ipsilateral CA1 regions, the ipsilateral dentate gyrus (DG), and the anterior cingulate cortex one month after injection in wild-type (WT) mice. Scale bars, 50 µm. **(E)** Quantification of MC1^+^ p-tau protein in different areas of brains from saline, non-phosphorylated tau, and p-tau injection groups. Shown are means ± SEM. P-vale was determined by one-way ANOVA with Tukey’s multiple comparison post-hoc test. (*P<0.05, ***p=0.0001, ****p<0.0001). **(F)** Schematic depiction of MC1 signals in the hippocampus and cortex.

P-tau causes apoptosis in cultured cells (*18, 20*). Given the potent neurotoxicity of p-tau *in vitro* (Fig. 1 and Sup Fig. 1), we posited that the spread of pathological tau would lead to loss of neuronal viability *in vivo*. To this end, we used NeuN to stain the nuclei of mature neurons, and used the TUNEL assay, which detected DNA fragmentation, to examine the fate of brain cells in hippocampus and AC cortex. Three months after p-tau injection, there was a significant reduction of NueN^+^ cells in the hippocampus and the AC cortex of the ipsilateral side after p-tau injection (arrows in Fig. 4A; middle and lower panels). In the meantime, elevation of TUNEL signals was observed (Fig. 4B-G), suggesting death of neurons in these regions. Among the NeuN^+^ cells in the p-tau group, many showed concomitant TUNEL signals, indicating that these neurons were on the path to death. Mice in other the tau or PBS control groups did not lose neurons at the ipsilateral side. The neuron loss was apparently an advanced trait of p-tau pathology, for one month after the injection there was no detectable difference in TUNEL staining among all groups of mice (Fig. 4D-G ).

**Fig. 4.**
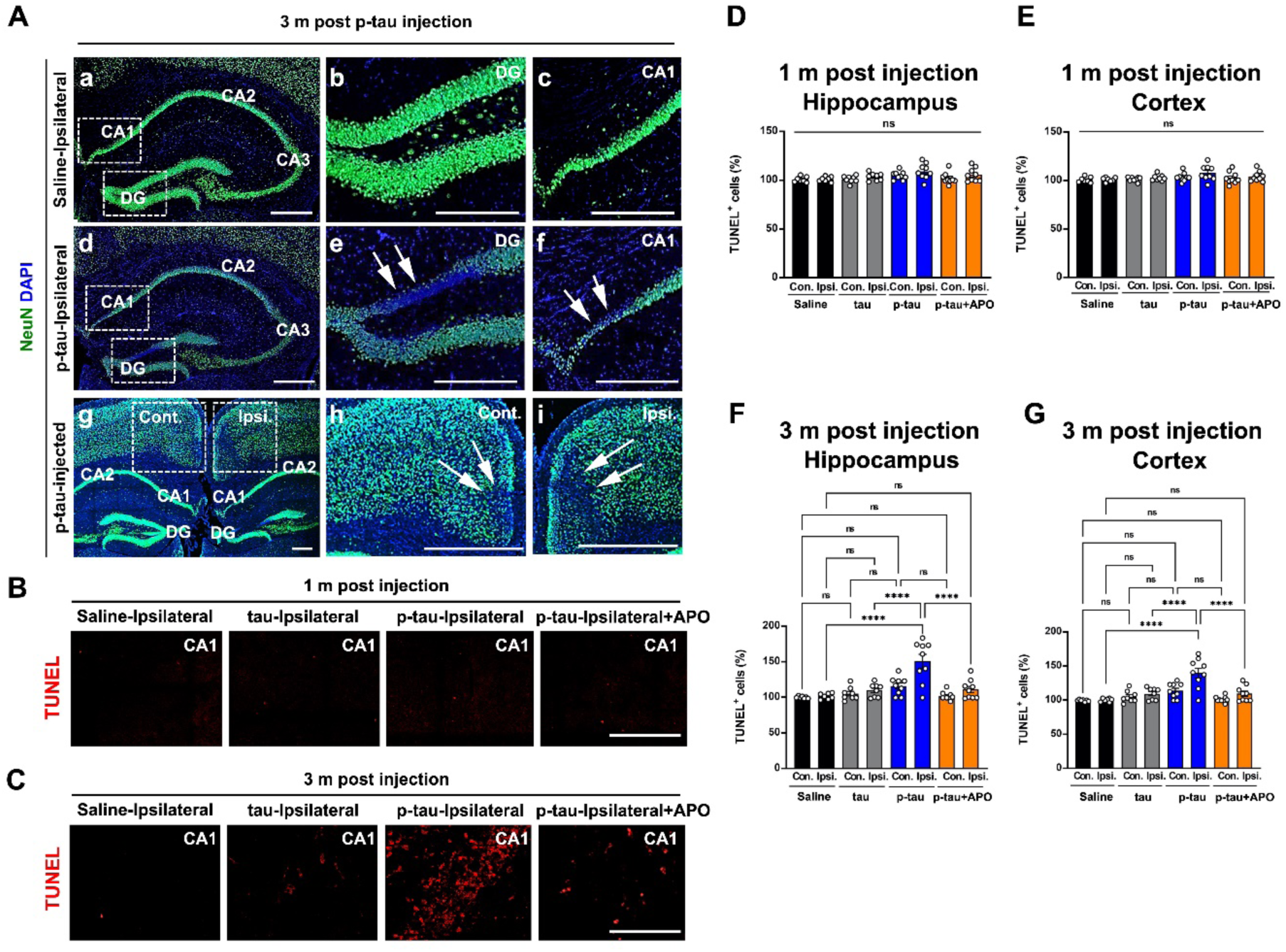
Apomorphine prophylactic treatment reduced neurotoxic effects following the injection of hyperphosphorylated tau into the hippocampus of WT mouse brains. **(A)** Intrahippocampal p-tau injection reduced neuron viability 3 months after injection. Shown are representative NeuN staining images of the dorsal hippocampus and anterior cingulate cortex in animals of the saline and p-tau groups. Neuronal loss was obvious in p-tau-injected hippocampus and anterior cingulate cortex. Panels b and c on the right are enlarged views of the squares indicated in the left panel a. Arrows in panels e and f highlight regions of the dentate gyrus (DG) and CA1 with a pronounced loss of NeuN signals. Similarly, panels h and i on the right are enlarged views of the squares shown in the left panel g, with arrows pointing to the contralateral and ipsilateral regions of the anterior cingulate cortex that exhibit significant NeuN signal loss. (B, C) The images depict TUNEL staining results in the ipsilateral CA1 sector under the specified conditions, one or three month post-injection. Scale bars, 50 µm. **(D, E, F, G)** Quantification of TUNEL^+^ cells in entire hippocampus and cortex in indicated groups. The number of TUNEL^+^ cells in saline-injected was set as 100%. Shown are means ± SEM. P-vale was determined by one-way ANOVA with Tukey’s multiple comparison post-hoc test. (****p<0.0001)

### P-tau injection causes time-dependent behavioral deficits

The spread of misfolded (i.e., the MC1 epitope, Fig. 3F) and hyperphosphorylated tau (i.e., AT8 epitope, Fig. 3C) from the hippocampus to the AC cortex, as well as the significant loss of neurons at the similar brain regions (Fig. 4) prompted us to assess whether neurological symptoms could result from p-tau injection. To this end, we conducted multiple behavioral tests, including the elevated plus maze (EPM) to assess anxiety (*26*), novel object recognition (NOR) examining short-term memory (*27*), and the Morris water maze test (MWM) for spatial memory (*28*). While the rotarod tests showed normal motor functions (Fig. 5D, E), p-tau-injected mice displayed significant aversion to open arms in the EPM tests in both 1- and 3-month post-injection animals (Fig. 5A-C), a result that suggested anxiety experienced by these animals. These results are consistent with the observations that anxiety is among the major neuropsychiatric symptoms predicting AD. Interestingly, the unphosphorylated tau also caused a moderate symptom 3 months after the injection. In the NOR test, mice with normal short-term memories show preference to explore a novel object over a familiar one. Inability to differentiate the familiar from a novel object indicates impaired short-term memory. Indeed, while mice that received PBS or tau were able to recognize the novel object, the p-tau group mice developed deficits in distinguishing the novel from the familiar object three months after the injection (Fig. 6A -C). The spatial memory tests of the Morris Water Maze further showed that, compared with the mice in the PBS and tau groups, p-tau group animals exhibited significantly longer escape latencies (Fig. 6D, G, H). Both memory tests revealed that the deficits were not noticeable 1-month post-injection (Fig. 6B, E, F). Together, these results show that injecting p-tau to the dorsal hippocampus caused behavioral and cognitive impairments in one to three months.

**Fig. 5.**
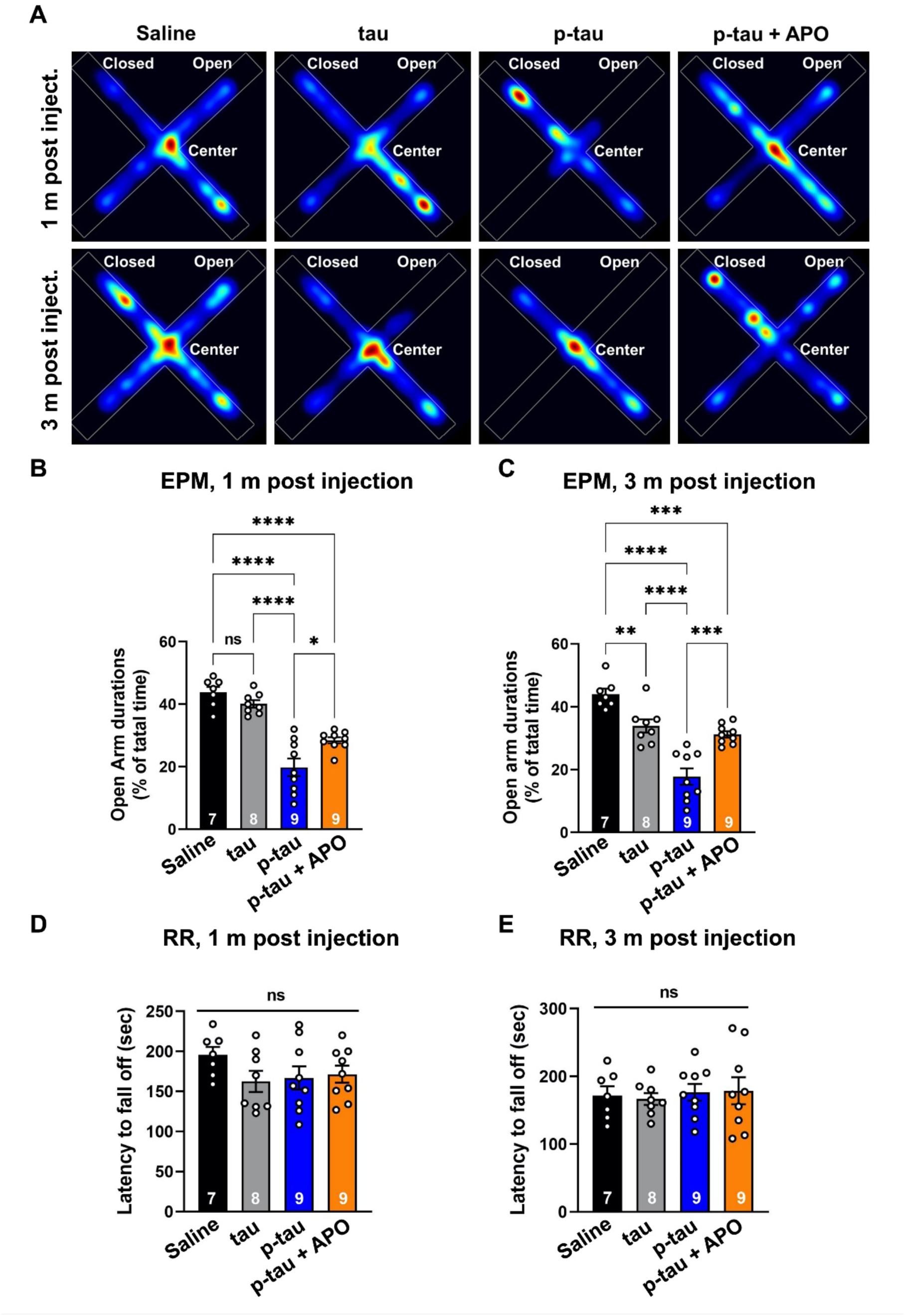
Intrahippocampal injection of hyperphosphorylated tau into the WT mouse brain induces anxiety-like behavior. **(A)** Stereotaxic injection of p-tau into the murine dorsal hippocampus induced aversion to open arms (a sign of anxiety) in WT male mice at 1m and 3m, as demonstrated by the Elevated Plus Maze (EPM) test. Prophylactic treatment with Apomorphine attenuated behavioral deficits. **(B, C)** Quantification results from A. Sample sizes for each group are labeled. Means ± SEM are presented (Saline, n=7 males; tau, n=8 males; p-tau, n=9 males; p-tau + APO, n=9 males). P-values were determined by one-way ANOVA with Tukey’s multiple comparison post-hoc test (*p<0.05, **p<0.01, ***p=0.0001, ****p<0.0001). **(D, E)** No differences were noted in the rotarod test in any group. RR, rotarod test. Shown are mean SEM. p-value was determined by one-way ANOVA with Tukey’s multiple comparison post-hoc test.

**Fig. 6.**
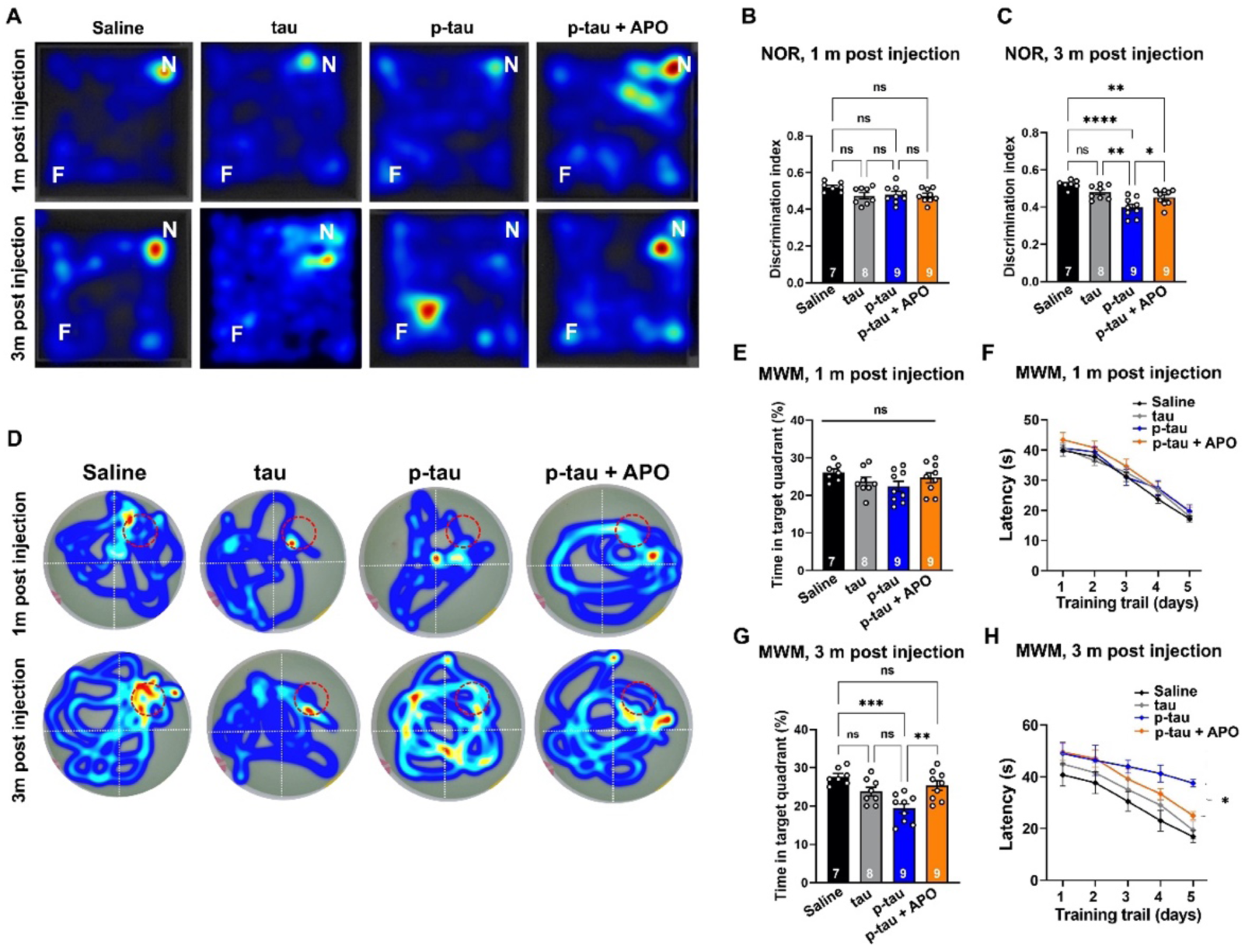
Prophylactic treatment of apomorphine mitigated cognitive deficits after hippocampal injection of hyperphosphorylated tau to WT mouse brain. (A-C) Intrahippocampal p-tau injection did not impair novel object recognition (NOR) at 1 month. However, at 3 months, intrahippocampal p-tau injection impaired NOR, an effect that was attenuated by APO treatment. Shown are the mean and SEM (Saline, n=7 males; tau, n=8 males; p-tau, n=9 males; p-tau + APO, n=9 males). The p-value was determined by one-way ANOVA with Tukey’s multiple comparison post-hoc test (*p<0.05, **p<0.01, ****p<0.0001). **(D-H)** The Morris water maze (MWM) test. Intrahippocampal p-tau injection impaired spatial learning memory in WT mice post 3 m, which was attenuated by the APO treatment. Probe trials were given on day 5 to evaluate retrieval performance of each group under various conditions. The p-tau group spent less time in the target platform than the saline, tau and p-tau + APO groups. Shown are the mean and SEM (Saline, n=7 males; tau, n=8 males; p-tau, n=9 males; p-tau + APO, n=9 males). The p-value was determined by one-way ANOVA with Tukey’s multiple comparison post-hoc test. (*P<0.05, **p<0.01, ***p=0.0001)

After establishing that p-tau caused neuron death and behavioral deficits in non-transgenic mice, we examined whether these pathological manifestations could be antagonized pharmacologically. Specifically, we tested the effects of apomorphine, which we found to afford highly effective protection against p-tau-inflicted cellular and subcellular damages (*18–20*). Apomorphine was administered to p-tau-injected mice via weekly subcutaneous injection at a dose of 10 mg/kg bodyweight, starting one week after the stereotactic p-tau injection. Animals were tested for the same panel of behavioral tests before sacrificing for immunohistochemical examination. Figures 5 and 6 showed that apomorphine effectively prevented the p-tau-induced anxiety and defects of short-term memory and spatial learning (Fig. 5 and 6). These benefits correlated with the maintenance of hippocampal and cortical neuronal integrity, as evidenced by the TUNEL assays (Fig. 4A, bottom row).

## DISCUSSION

In summary, we show that hyperphosphorylated tau confers neurotoxicity that is consistent with the hypothesis that neurodegeneration of AD and other tauopathies results from the propagation of pathological tau-inflicted damages in the brain. By injecting 2 µg of p-tau oligomers to the hippocampus, pathologically relevant tau features, such as the AT8 and MC1 epitopes, spread from the site of injection to distal cortical regions in three months. Prominent signs of neuronal loss and apoptosis ensue. These pathological manifestations are accompanied by defects in behaviors and cognition. Equally important is the demonstration that these cellular and neurological abnormalities are effectively prevented by subcutaneous injection of apomorphine, thereby paving the way for using this model to support screens and tests of compounds for AD therapeutics.

This work presents evidence that hyperphosphorylated tau oligomers cause neurodegenerative traits in non-transgenic wildtype mice. Many widely used animal models for AD either lack the element of tau hyperphosphorylation, or rely on the transgenic expression, frequently overexpression, of a mutant tau associated with certain types of frontotemporal dementia, but not AD per se. Intrahippocampal injection of tau extracted from AD or other tauopathy brain specimens faithfully recapitulate the original tau pathologies (*12–15*), hence complying with the Koch’s postulates for identifying the appropriate pathogen vis-à-vis neurofibrillary tangles and related tau deposits. However, no significant cognitive deficits were seen in these mice, except when a human mutation(s) in tau or the Aβ biogenesis pathway is already present. Even the humanized mice with the rodent tau gene completely replaced by the human orthologue maintained cognitive integrity after receiving the AD tau extract injection to the hippocampus (*16*). These results raise the question as to whether the disease-causing species of tau is appropriately presented in these brain extracts that favor the formation of the conspicuous tangles. The lack of neurological deficits in these mice is consistent with the findings that tau tangles in neurons impose negligible effects on the function and viability of the underlying neurons (*29*). Transgenic flies expressing human tau in neurons show neurological defects without detectable neurofibrillary tangles (*30*).

It is worth noting that the p-tau used in this work was retrieved from freezer stocks stored immediately after bacterial production and the subsequent purification process. No pre-incubation for fibrillogenesis before injection was needed. P-tau was isolated via its exceptional thermotolerance while other proteins in the bacterial lysate were precipitated in a boiling water bath. When applied to size exclusion chromatography, the majority of p-tau from the PIMAX expression system was eluted as oligomers ranging from 66 to 400 kDa (Fig. 1A), corresponding to 2- to 10-mers. Repetitive attempts of transmission electron microscopy failed to reveal a predominant, uniform conformation (*18, 20*), an anticipated scenario from the highly intrinsically disordered nature of tau as predicted by AlphaFold or other algorithms (not shown). Due either to the thermotolerance or to refolding during the purification after heat treatment, certain p-tau molecules maintain a quaternary structure that confers neurotoxicity and nucleates the aggregation and cytotoxicity of unmodified tau (*18*). Apparently, a window exists for small-molecule compounds to bind and to modulate the cytotoxic tau conformers. Among the most notable are apomorphine and raloxifene that antagonize, whereas a group of benzodiazepines enhance p-tau aggregation and cytotoxicity in vitro (*19*). Given the well-documented functional and physical association between pathological tau and other biomolecules including proteins (*25, 31*), lipids (*32*), glycans (*33*), and RNAs (*34*), it is tempting to speculate that the transformation of hyperphosphorylated tau molecules to a disease-triggering species, as opposed to the thermodynamically favorable neurofibrillary tangles, is determined by stochastic encountering with metabolites or biomolecules in the brain that are deemed as risk factors of AD.

Apomorphine was found to be as a potent p-tau aggregation and cytotoxicity inhibitor in a pilot screen of a small prescription drug library (*19*). This compound blocks essentially all the ill-effects imposed by p-tau to SH-SY5Y cells including ER stress, mitochondrial dysfunction, and apoptosis (*20*). In addition to the wildtype p-tau, apomorphine also neutralizes the detrimental effects of p-tau bearing the P301L mutation that is linked to frontotemporal dementia (*35*). The benefits of this drug therefore are likely to extend to other tauopathies such as FTD and chronic traumatic encephalopathies. Besides apomorphine, the same screen identified another prescription drug, raloxifene, for a comparable anti-p-tau potency. Raloxifene reduces the risk of dementia in women (*36*) but has no clear benefits to those already showing mild to moderate severity of Alzheimer’s disease (*37*). Experiments presented in this work were designed to test the feasibility of a new mouse model for hyperphosphorylated tau-driven neuropathology. The drug administration regimen verifies the prophylactic potential of apomorphine. Whether giving this drug at a later time could afford therapeutic efficacy remains to be examined. For its dopamine receptor agonist activity (*38*), apomorphine is currently used to treat the off episodes in Parkinson’s disease patients. The inconvenience of subcutaneous injection and the need of an anti-emetic medicine to reduce a side-effect of apomorphine present a challenge for the use of apomorphine to treat AD. Nonetheless, apomorphine is the oldest antiparkinsonian drug on the market (*39*), and new formulations supporting different delivery strategies are available (*40*). It will be interesting to survey whether regular users of apomorphine have lower prevalence of Alzheimer’s disease or related dementia.

## Materials and Methods

### Preparation of p-tau

The hyperphosphorylated tau (1N4R isoform) was induced and purified according to previously published^17^. To examine the size of the p-tau to be subjected to mouse injection, size exclusion chromatography on a GE Healthcare Amersham AKTA FPLC was performed. The TEV protease-treated p-tau sample (0.5 ml) was loaded to a Superdex 200 10/300 GL size exclusion column, and resolved by passing 20 mM Tris pH 7.4 through the column at a flow rate of 0.5 ml/min. 1-ml fractions were collected and examined by SDS-PAGE. The fractions were compared to standards thyroglobulin (669kD), ferritin (440 kD), BSA (66 kD), and Ribonuclease A (14 kD). P-tau-containing fractions were pooled and concentrated by centrifugation through the Amicon 30 kD spin column at 5,000 x g for 20 minutes until the volume dropped to below 2 ml. 4 ml of purification buffer (20 mM Tris pH7.4, 100 mM NaCl) was added to the spin column, mixed, and resumed centrifugation. The final solution with p-tau was collected and mixed with final 10% (v/v) glycerol for storage at -80°C until use.

### Primary neuron culture

Mouse cortical neurons were isolated from E16.5 mouse embryos. The cortex was mechanically dissociated by passing it through a glass pipette twenty times and then filtered through a 70-µm nylon mesh filter (BD Biosciences). 2 X 10^5^ cells were plated onto plastic culture plates coated with 30 µg/ml poly-L-lysine and 2 µg/ml laminin, and cultured in Neurobasal Media supplemented with 10% FBS, 100 U/ml penicillin and streptomycin, and N2 and B27 supplements (Invitrogen). The cultures were maintained at 37°C in a humidified atmosphere with 5% CO2 and 95% air. At 6 days *in vitro* (D.I.V.), recombinant tau or p-Tau proteins were added to the culture medium, respectively.

### MitoTracker labeling and MitoSOX assay

Primary mouse cortical neurons were plated on 35 mm Glass bottom dish (MatTek Life Science). Neurons were cultured to 6 DIV, and then exposed to recombinant tau and p-Tau proteins 2 day for MitoSOX imaging. At 8 DIV, the cells are labeled by the MitoSOX™ Red Mitochondrial Superoxide Indicators (Invitrogen) or MitoTracker® Green FM (Invitrogen) according to the manufacturer’s instructions.

### Protein extraction and western blotting

Primary mouse cortical neurons were washed once with PBS and dissolved in radioimmunoprecipitation assay lysis buffer (20 mm HEPES, pH 7.8, 150 mm NaCl, 1 mm EDTA, 0.1% Triton X-100, 50 mm NaF, 1 mm dithiothreitol, and protease inhibitor cocktail). The cell lysates were cleared by centrifugation at 13,000 × g for 10 min, and the supernatants were used for Western blotting analysis. The protein concentration was determined using a BCA kit (Pierce). Equal amounts of either tissue or cell lysates were subjected to SDS-PAGE and transferred to polyvinylidene difluoride membranes. The membrane was blocked with 5% skimmed milk and probed with the primary antibody at 4°C overnight. After the membrane was washed and incubated with HRP-conjugated secondary antibodies at room temperature for 1 h, it was developed with an ECL kit (GE Healthcare). The following antibodies were used: rabbit anti-Map2 (Abcam); rabbit anti α -Tubulin (Cell Signaling Technology); mouse anti TuJ1 (Covance); rabbit anti-Cleaved Caspase 3 (Cell Signaling Technology); mouse anti Phospho-Tau (Ser202, Thr205) (AT8) (Thermo Fisher Scientific); mouse anti Flag M2 (Sigma); rabbit anti-GAPDH (Cell Signaling Technology).

### Propidium Iodide assay

Propidium iodide (1.0 mg/mL solution in water) (ThermoFisher) was incubated with 10 D.I.V. cortical neurons for 1 hr at a concentration of 6 µM. During imaging PI was typically excited at 800 nm, but where indicated was excited off-peak so as to excite another fluorophore simultaneously for co-registration. PI emission was detected through the 570-670 nm filter. The stained suspensions were collected with flow cytometric analyses (Attune, Applied Biosystems) and analyzed with FlowJo v10.

### Animals and stereotaxic surgery

After deeply anesthetizing 2- to 3-month-old WT male mice using a ketamine/xylazine mixture, the animals were immobilized in a stereotaxic frame (Stoelting Co). Stereotaxic injections were then performed using a WPI nanoliter 2020 injector coupled with pre-pulled glass pipettes to deliver the drug into the hippocampus (coordinates: bregma, −2.5 mm; lateral, +2 mm; depth, −1.8 mm). All injected animals were monitored during and after the surgery, with analgesics administered postoperatively. All experiments adhered to protocols approved by the University of Virginia Animal Care and the National Yang Ming Chiao Tung University Animal Care and Use Committee, in compliance with the guidelines of the Association for Assessment and Accreditation of Laboratory Animal Care.

### Immunohistochemistry

Brains from the indicated conditions were fixed in 4% paraformaldehyde (PFA) at 4°C overnight with gentle agitation, then cryopreserved in 30% sucrose, frozen, and stored at −20°C until use. For staining, 20-µm cryosections were prepared and incubated in a blocking/permeabilization solution containing 3% normal goat serum (NGS) and 0.2% Triton-X in PBS. The sections were treated overnight with appropriate primary antibodies diluted in 1% NGS/0.2% Triton-X100/PBS, followed by incubation with the corresponding secondary antibodies for 2 hours at room temperature. The primary antibodies used included: mouse anti-phospho-tau (Ser202, Thr205) (AT8) (Invitrogen); mouse anti-MC1 (from Dr. Peter Davies)(*24*); rabbit anti-IbaI (Wako Chemicals USA Inc.); rabbit anti-NeuN (Abcam); and mouse anti-tau (DA9) (a gift by P. Davies).

### TUNEL assay

Brains from the indicated littermates were fixed in 4% paraformaldehyde at 4°C overnight with gentle agitation, then cryopreserved in 30% sucrose, frozen, and stored at −20°C until the TUNEL assay was performed. TUNEL staining was used to identify apoptotic cells under light microscopy using a Click-iT TUNEL Alexa Fluor 594 Imaging Assay kit (Thermo Fisher Scientific), following the manufacturer’s instructions.

### Apomorphine injection

Apomorphine (Cayman Chemical, Ann Arbor, MI; Cat. #16094) was dissolved in saline and administered subcutaneously once a week, 5x, at a concentration of 10 mg/kg, based on previous studies involving apomorphine injection in Alzheimer’s disease model mice (20). Saline-only injections were used as a control.

### Elevated plus maze (EPM) test

Mice were placed in the center of an elevated plus maze (arms 6.2 × 75 cm, with side walls 20 cm high on the two closed arms, elevated 63 cm above the ground) for 5 minutes to assess anxiety. Automated scoring was used to measure the cumulative time each mouse spent in the open and closed arms of the maze.

### Novel object recognition (NOR) test

Mice naturally display a preference for novel objects. The experimental apparatus was a 25 × 25 × 25 cm box made of acrylic plates, placed in a separate room. Videos of the animals’ behavior were captured and analyzed. The experiment consisted of three phases: habituation, training, and testing. During the training phase, mice were allowed to explore two identical objects for 10 minutes. One hour later, one of the familiar objects was replaced with a novel object for testing. If the mouse recognizes the familiar object, it will spend more time with the novel object, indicating normal recognition memory. The recognition index for the novel object was calculated to assess the effect on recognition memory.

### Rotarod

Motor performance and motor learning were assessed using a rotarod apparatus (Ugo Basile, Germany). Each mouse was weighed before testing. The rotarod has six lanes, allowing six mice to be tested simultaneously. Each group of six mice underwent five trials, with a maximum duration of 5 minutes (300 seconds) per trial, the rotarod was accelerated from 5 rpm to 40 rpm in 300s. The latency to fall of each rat was recorded. The latency to fall was recorded as a measure of motor coordination.

### Morris Water Maze Analysis

The Morris Water Maze (MWM) test was conducted, following a protocol adapted from Vorhees (*28*). Briefly, a submerged platform was placed in the center of one quadrant of a round swimming pool, with four conspicuous spatial cues surrounding the pool. During the training phase, each mouse underwent four trials per day, with a 10 to 20-minute interval between trials. If a mouse failed to locate the platform within 60 seconds, it was manually guided to the platform and allowed to remain there for 10 seconds. An overhead camera recorded the swimming paths. The day after training, the mice were released from the opposite quadrant, and their movements were recorded. Video recordings were analyzed using Noldus EthoVision XT software to assess the animals’ movement patterns and time spent in each quadrant. The performance data from all trials of each group were pooled for analysis.

### Quantification of MAP2 fluorescence

Images were acquired using a SP8 confocal microscope with 20 magnification with an image matrix of 1024 1024pixel, a pixel scaling of 0.2 x 0.2 mm and a depth of 8 bit. Confocal images were collected in z stacks with a slice distance of 0.4 mm. Analyses of immunostaining and cell quantification were performed using the National Institutes of Health ImageJ software. for MAP2 analysis. Specifically, the intensity of the background and the signal was obtained by determining the background and signal threshold levels using the Threshold tool across at least three different fields corresponding to the differentiation step with the highest signal intensity. The average threshold levels were then calculated and applied to all images under analysis using the ’Set Threshold’ tool. After setting the thresholds, the MFI of the background and the signal was obtained using the Measure tool, with the background intensity subtracted from the signal intensity. This analysis was performed on at least three fields for each differentiation step. The resulting values were then averaged and subjected to statistical analysis.

### Statistical analysis

Data were summarized as mean ± SEM. Statistical tests were conducted independently using Prism 7 and R 3.6.0 to ensure consistency, with specific tests indicated in the figures. Briefly, one-way analysis of variance (ANOVA) followed by Tukey’s post hoc test for multiple comparisons was used to compare three or more groups. A P value of less than 0.05 was considered statistically significant. Power analysis and sample size determination were performed in R with a significance level of 0.05 and 80% statistical power. Figures were generated using Prism 7.

## Acknowledgments

**Funding:** This work was supported by NIH grants (NS135693 to C.-Y.K. and M.-H.K; AG062435, AG077475, AG057274 to M.-H.K.; NS125677, NS125788, HD109025 to C.- Y.K.), NSTC grants (NSTC111-2320-B-A49-043 and NSTC112-2628-B-A49-011-MY3 to H.-R.C.), Yushan Fellow Program and Brain Research Center, National Yang Ming Chiao Tung University from The Featured Areas Research Center Program within the framework of the Higher Education Sprout Project by the Ministry of Education (MOE) in Taiwan to H.-R.C.

**Author contributions:** H.-R.C., M.-H.K. and C.-Y.K. designed the study and wrote the manuscript. H.-R.C., H.-T. H., K.-W.W., S.H., H.-Y.L., M.-H.L., C.-C.C, C.-Y.W., C.-Y.C., E. B., and M. K. conducted the experiments. H.-R.C. performed the statistical analysis.

**Competing interests:** The authors declare that they have no competing interests.

**Data and materials availability:** All data needed to evaluate the conclusions in the paper are present in the paper and/or the Supplementary Materials. Additional data related to this paper may be requested from the authors.

---

**Supplemental Fig. 1.**
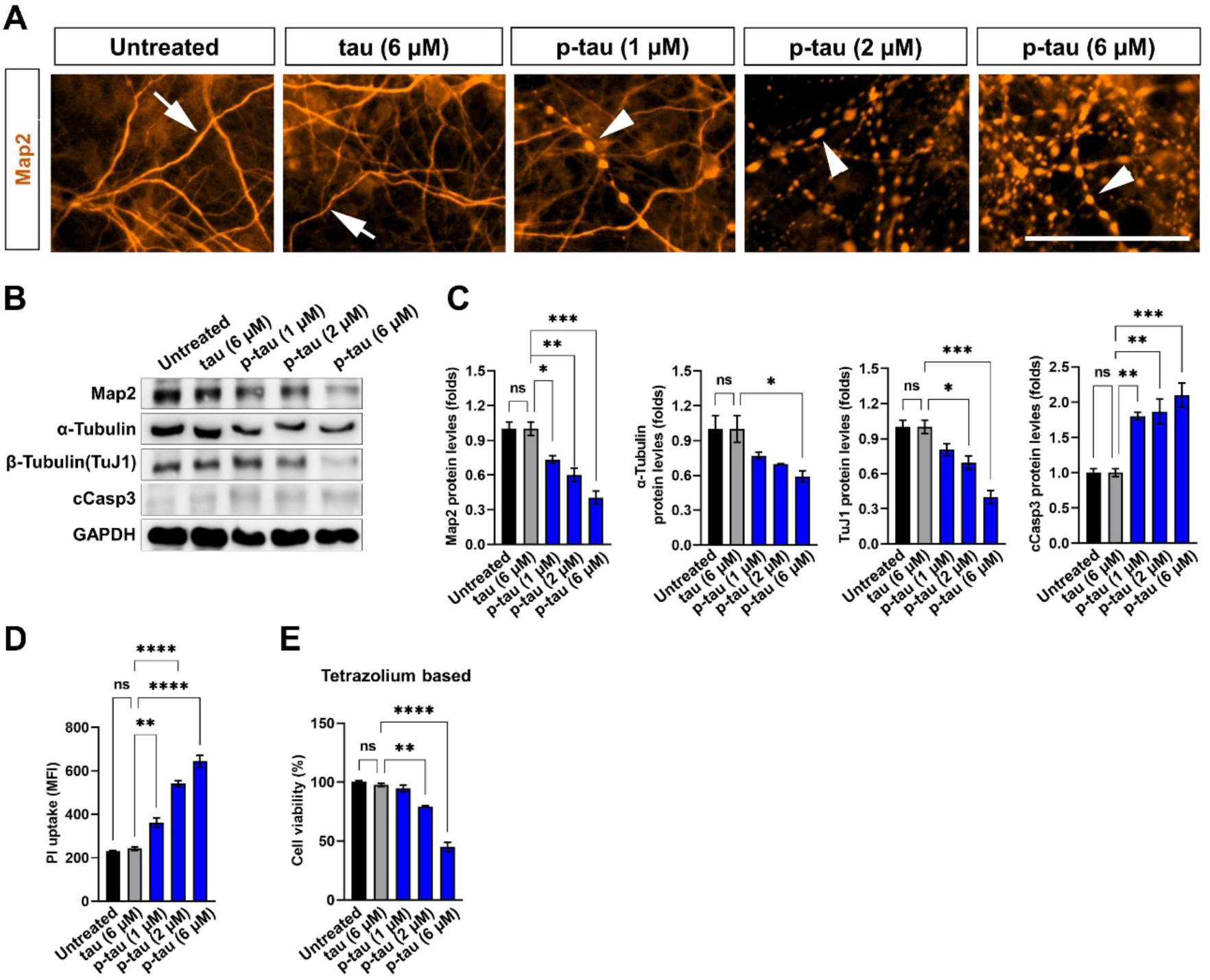
Hyperphosphorylated human tau oligomers induces neurite fragmentation while non-phosphorylated tau exhibits no such effects. Cortical neurons isolated from embryonic day 16.5 (E16.5) C57BL/6 mice were cultured in vitro for 6 days, followed by a 2- or 4-day treatment with varying concentrations of non-phosphorylated tau or p-tau oligomers. (A) Neurites of cortical neurons showed p-tau dose-dependent fragmentation. Neurites were visualized by immunofluorescence staining with anti-Map2 antibodies on neurons treated with p-tau or tau for 4 days. Clear fragmentation is seen in p-tau, but not in tau-treated neurons (see arrows). The arrows indicate normal neurites, while the arrowheads point to fragmented neurites. Scale bars, 50 µm. (B) P-tau caused deficiencies of proteins key to neurites development. Neuron lysates (4-day treatment) were resolved and probed with anti-Map2, anti-α tubulin, anti-TuJ1 and anti-cleaved Caspase 3 antibodies. The representative images of the immunoblotting are shown in panel (B). Panel (C) shows the quantification results. Data are means ± SEM (n = 6), and statistical analysis was performed using one-way ANOVA with Tukey’s post hoc test. (*P<0.05, **p<0.01, ***p=0.0001) (D, E) Propidium Iodide uptake and tetrazolium-based viability assays showed the neurotoxicity caused by ptau at 4 days. Shown are the mean and SEM from 3 sets of independent cultures. (**p<0.01, ****p<0.0001)

**Supplemental Fig. 2.**
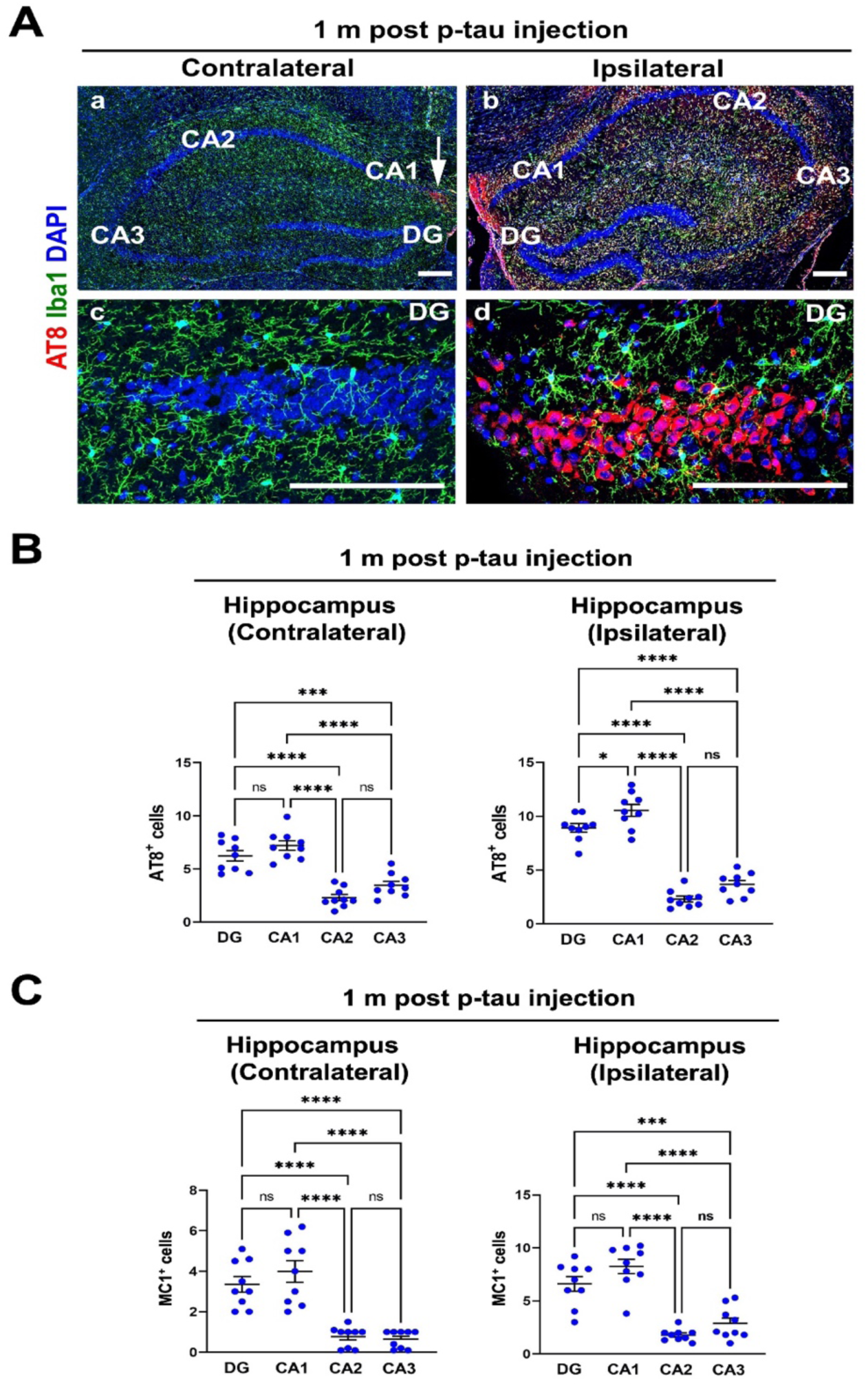
Engulfment of aggregated p-Tau (AT8+ neurons) by microglia and macrophages following hippocampal injection of hyperphosphorylated tau. **(A)** One month after the p-tau injection, numerous AT8^+^ cells accumulated in the dentate gyrus (DG) of the ipsilateral hippocampus, surrounded by Iba1+ activated microglia in panel d. On the contralateral side, AT8 signals were observed in the fimbria area, as indicated by the arrow in panel a. Scale bars, 50 µm. **(B, C)** Quantification of AT8^+^ p-tau protein in different areas of brains from saline, non-phosphorylated tau, and p-tau injection groups. Shown are means ± SEM. P-vale was determined by one-way ANOVA with Tukey’s multiple comparison post-hoc test. (*P<0.05, ***p=0.0001, ****p<0.0001).

**Supplemental Fig. 3.**
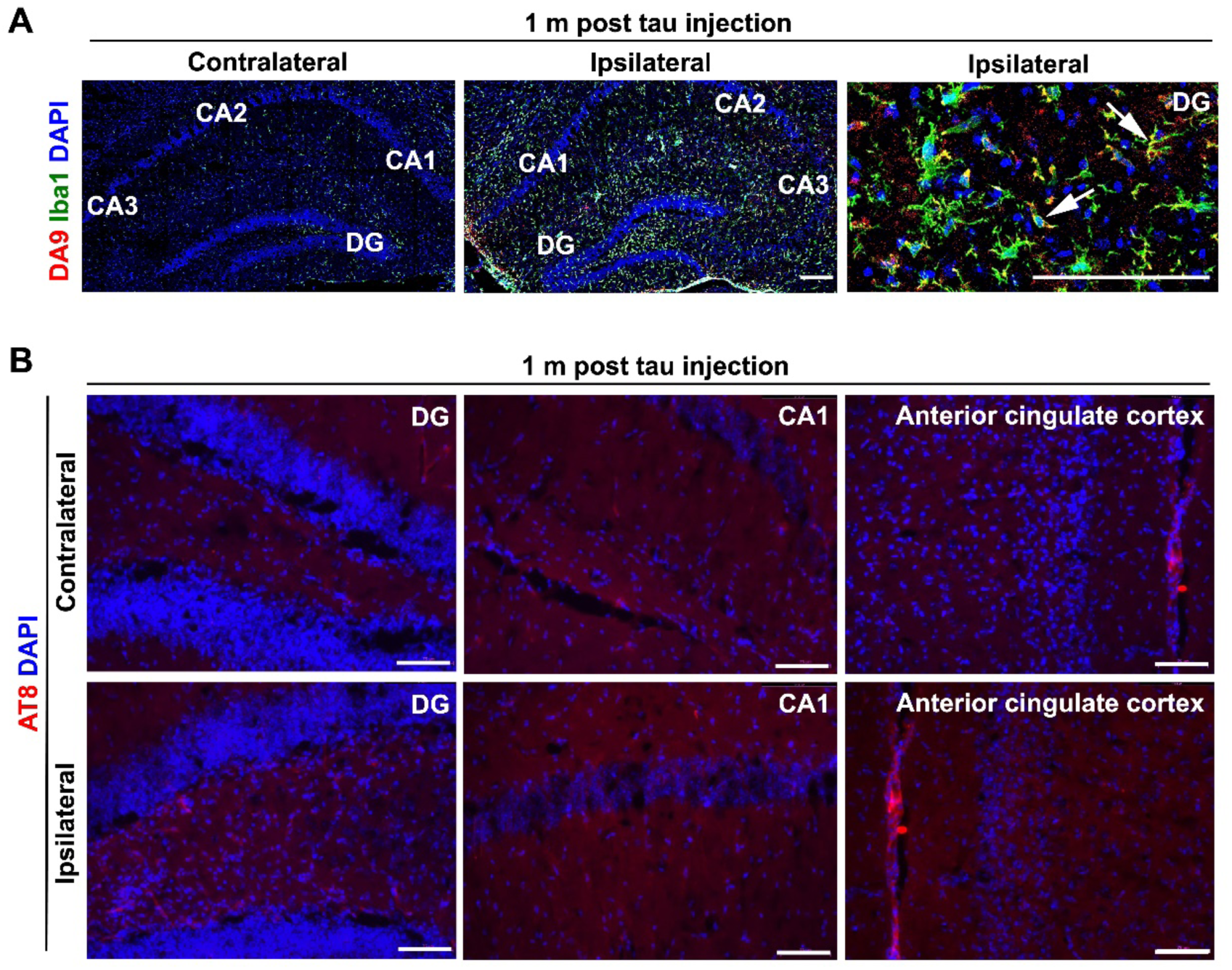
Mice injected with non-phosphorylated tau did not exhibit AT8 signals in either the hippocampus or cortex one month post-injection. (A) DA9^+^ pan-tau staining at 1m post non-phosphorylated tau injection. Almost all recombinant tau was engulfed by Iba1^+^ microglia post-injection (arrows). Scale bars, 50 µm. (B) Tau-injected mice did not show AT8 signals in both the hippocampus and cortex at 1 month post-injection. Scale bars, 50 µm.

## References

1. I. Grundke-Iqbal, et al., Microtubule-associated protein tau. A component of Alzheimer paired helical filaments. J Biol Chem 261, 6084–6089 (1986).

2. I. Grundke-Iqbal et al., Abnormal phosphorylation of the microtubule-associated protein tau (tau) in Alzheimer cytoskeletal pathology. Proc Natl Acad Sci U S A 83, 4913–4917 (1986).

3. H. Braak, E. Braak, Neuropathological stageing of Alzheimer-related changes. Acta Neuropathol 82, 239–259 (1991).

4. E. D. Roberson et al., Reducing endogenous tau ameliorates amyloid beta-induced deficits in an Alzheimer’s disease mouse model. Science 316, 750–754 (2007).

5. J. M. Nussbaum et al., Prion-like behaviour and tau-dependent cytotoxicity of pyroglutamylated amyloid-beta. Nature 485, 651–655 (2012).

6. M. Goedert, M. G. Spillantini, Tau mutations in frontotemporal dementia FTDP-17 and their relevance for Alzheimer’s disease. Biochim Biophys Acta 1502, 110–121 (2000).

7. G. G. Kovacs, Invited review: Neuropathology of tauopathies: principles and practice. Neuropathol Appl Neurobiol 41, 3–23 (2015).

8. Y. Soeda, A. Takashima, New Insights Into Drug Discovery Targeting Tau Protein. Front Mol Neurosci 13, 590896 (2020).

9. X. Y. Sun et al., Fc effector of anti-Abeta antibody induces synapse loss and cognitive deficits in Alzheimer’s disease-like mouse model. Signal Transduct Target Ther 8, 30 (2023).

10. N. J. Reish et al., Multiple Cerebral Hemorrhages in a Patient Receiving Lecanemab and Treated with t-PA for Stroke. N Engl J Med 388, 478–479 (2023).

11. F. M. LaFerla, K. N. Green, Animal models of Alzheimer disease. Cold Spring Harb Perspect Med 2, (2012).

12. Z. He et al., Transmission of tauopathy strains is independent of their isoform composition. Nat Commun 11, 7 (2020).

13. J. L. Guo et al., Unique pathological tau conformers from Alzheimer’s brains transmit tau pathology in nontransgenic mice. J Exp Med 213, 2635–2654 (2016).

14. S. G. Greenberg, P. Davies, A preparation of Alzheimer paired helical filaments that displays distinct tau proteins by polyacrylamide gel electrophoresis. Proc Natl Acad Sci U S A 87, 5827–5831 (1990).

15. F. Clavaguera et al., Brain homogenates from human tauopathies induce tau inclusions in mouse brain. Proc Natl Acad Sci U S A 110, 9535–9540 (2013).

16. T. Saito et al., Humanization of the entire murine Mapt gene provides a murine model of pathological human tau propagation. J Biol Chem 294, 12754–12765 (2019).

17. D. Sui et al., Protein interaction module-assisted function X (PIMAX) approach to producing challenging proteins including hyperphosphorylated tau and active CDK5/p25 kinase complex. Mol Cell Proteomics 14, 251–262 (2015).

18. M. Liu et al., Hyperphosphorylation Renders Tau Prone to Aggregate and to Cause Cell Death. Mol Neurobiol 57, 4704–4719 (2020).

19. M. Liu et al., Hyperphosphorylated tau aggregation and cytotoxicity modulators screen identified prescription drugs linked to Alzheimer’s disease and cognitive functions. Sci Rep 10, 16551 (2020).

20. Z. Song et al., Hyperphosphorylated Tau Inflicts Intracellular Stress Responses that Are Mitigated by Apomorphine. Mol Neurobiol 61, 2653–2671 (2024).

21. E. Himeno et al., Apomorphine treatment in Alzheimer mice promoting amyloid-beta degradation. Ann Neurol 69, 248–256 (2011).

22. M. Iba et al., Synthetic tau fibrils mediate transmission of neurofibrillary tangles in a transgenic mouse model of Alzheimer’s-like tauopathy. J Neurosci 33, 1024–1037 (2013).

23. J. Biernat et al., The switch of tau protein to an Alzheimer-like state includes the phosphorylation of two serine-proline motifs upstream of the microtubule binding region. EMBO J 11, 1593–1597 (1992).

24. G. A. Jicha, R. Bowser, I. G. Kazam, P. Davies, Alz-50 and MC-1, a new monoclonal antibody raised to paired helical filaments, recognize conformational epitopes on recombinant tau. J Neurosci Res 48, 128–132 (1997).

25. J. J. Siew et al., Galectin-3 aggravates microglial activation and tau transmission in tauopathy. J Clin Invest 134, (2024).

26. A. A. Walf, C. A. Frye, The use of the elevated plus maze as an assay of anxiety-related behavior in rodents. Nat Protoc 2, 322–328 (2007).

27. M. Leger et al., Object recognition test in mice. Nat Protoc 8, 2531–2537 (2013).

28. C. V. Vorhees, M. T. Williams, Morris water maze: procedures for assessing spatial and related forms of learning and memory. Nat Protoc 1, 848–858 (2006).

29. K. V. Kuchibhotla et al., Neurofibrillary tangle-bearing neurons are functionally integrated in cortical circuits in vivo. Proc Natl Acad Sci U S A 111, 510–514 (2014).

30. C. W. Wittmann et al., Tauopathy in Drosophila: neurodegeneration without neurofibrillary tangles. Science 293, 711–714 (2001).

31. J. N. Rauch et al., LRP1 is a master regulator of tau uptake and spread. Nature 580, 381–385 (2020).

32. D. Toral-Rios et al., Cholesterol 25-hydroxylase mediates neuroinflammation and neurodegeneration in a mouse model of tauopathy. J Exp Med 221, (2024).

33. J. Zhao et al., 3-O-Sulfation of Heparan Sulfate Enhances Tau Interaction and Cellular Uptake. Angew Chem Int Ed Engl 59, 1818–1827 (2020).

34. L. Jiang et al., Interaction of tau with HNRNPA2B1 and N(6)-methyladenosine RNA mediates the progression of tauopathy. Mol Cell 81, 4209–4227 e4212 (2021).

35. K. W. Wang, G. Zhang, M. H. Kuo, Frontotemporal Dementia P301L Mutation Potentiates but Is Not Sufficient to Cause the Formation of Cytotoxic Fibrils of Tau. Int J Mol Sci 24, (2023).

36. K. Yaffe et al., Effect of raloxifene on prevention of dementia and cognitive impairment in older women: the Multiple Outcomes of Raloxifene Evaluation (MORE) randomized trial. Am J Psychiatry 162, 683–690 (2005).

37. V. W. Henderson et al., Raloxifene for women with Alzheimer disease: A randomized controlled pilot trial. Neurology 85, 1937–1944 (2015).

38. J. A. Subramony, Apomorphine in dopaminergic therapy. Mol Pharm 3, 380–385 (2006).

39. M. Auffret, S. Drapier, M. Verin, The Many Faces of Apomorphine: Lessons from the Past and Challenges for the Future. Drugs R D 18, 91–107 (2018).

40. F. Carbone, A. Djamshidian, K. Seppi, W. Poewe, Apomorphine for Parkinson’s Disease: Efficacy and Safety of Current and New Formulations. CNS Drugs 33, 905–918 (2019).

